# Hepatitis C virus infection is inhibited by a non-canonical antiviral signaling pathway targeted by NS3-NS4A

**DOI:** 10.1101/625640

**Authors:** Christine Vazquez, Chin Yee Tan, Stacy M. Horner

## Abstract

The hepatitis C virus (HCV) NS3-NS4A protease complex is required for viral replication and is the major viral innate immune evasion factor. NS3-NS4A evades antiviral innate immunity by inactivating several proteins, including MAVS, the signaling adaptor for RIG-I and MDA5, and Riplet, an E3 ubiquitin ligase that activates RIG-I. Here, we identified a Tyr-16-Phe (Y16F) change in the NS4A transmembrane domain that prevents NS3-NS4A targeting of Riplet but not MAVS. This Y16F substitution reduces HCV replication in Huh7 cells, but not in Huh-7.5 cells, known to lack RIG-I signaling. Surprisingly, deletion of RIG-I in Huh7 cells did not restore Y16F viral replication. Rather, we found that Huh-7.5 cells lack Riplet expression and that addition of Riplet to these cells reduced HCV Y16F replication. In addition, IRF3 deletion in Huh7 cells was sufficient to restore HCV Y16F replication, and the Y16F protease lacked the ability to prevent IRF3 activation or interferon induction. Taken together, these data reveal that the NS4A Y16 residue regulates a non-canonical Riplet-IRF3-dependent, but RIG-I-MAVS-independent, signaling pathway that limits HCV infection.

**Importance:** The HCV NS3-NS4A protease complex facilitates viral replication by cleaving and inactivating the antiviral innate immune signaling proteins MAVS and Riplet, which are essential for RIG-I activation. NS3-NS4A therefore prevents IRF3 activation and interferon induction during HCV infection. Here, we uncover an amino acid residue within the NS4A transmembrane domain that is essential for inactivation of Riplet, but does not affect MAVS cleavage by NS3-NS4A. Our study reveals that Riplet is involved in a RIG-I- and MAVS-independent signaling pathway that activates IRF3 and that this pathway is normally inactivated by NS3-NS4A during HCV infection. Our study selectively uncouples these distinct regulatory mechanisms within NS3-NS4A and defines a new role for Riplet in the antiviral response to HCV. As Riplet is known to be inhibited by other RNA viruses, such as such influenza A virus, this innate immune signaling pathway may also be important in controlling other RNA virus infections.

## Introduction

Hepatitis C virus (HCV) is a positive-sense, singled-stranded RNA virus that infects over 70 million people worldwide, with up to 80% of infected individuals developing chronic infection (1). The recent development of direct-acting antivirals for HCV has dramatically improved successful treatment of HCV infection (2). However, many HCV-infected individuals are asymptomatic and thus unaware of their HCV status until secondary manifestations, such as liver cirrhosis and hepatocellular carcinoma, arise decades later. Notably, although the current direct-acting antivirals treat HCV-induced disease, they do not always prevent re-infection in cured individuals. Therefore, there is an urgent need for future studies into the development of a vaccine to reduce the global burden of HCV infection (3).

Several factors contribute to the ability of HCV to establish a chronic infection, including its ability to evade detection and dysregulate the host antiviral innate immune response through the actions of the HCV NS3-NS4A protease complex (4). The NS3-NS4A protease is a protein complex formed between NS3, which contains protease and helicase domains, and NS4A. NS4A is a 54 amino acid protein that contains an N-terminal transmembrane domain, an NS3 interacting domain, and a C-terminal acidic domain (5). The NS4A transmembrane domain anchors NS3 to membranes (6) and mediates NS4A dimerization (7). NS3-NS4A has diverse functions in the HCV life cycle, with roles in HCV RNA replication, viral assembly, and innate immune evasion (reviewed in (8))(9). The mechanisms that regulate these diverse functions of NS3-NS4A are not completely understood. However, it is known that NS4A directs the protease complex to distinct intracellular membranes to perform some of these functions: the ER for viral replication; and mitochondria and mitochondrial-ER contact sites (often referred to as mitochondrial-associated ER membranes (MAM)) for immune evasion (10–14).

Antiviral innate immune signaling against HCV can be initiated by the RNA sensor proteins RIG-I and MDA5 (15–17). RIG-I is directly activated by multiple ubiquitination events by E3 ubiquitin ligases, namely TRIM25 and Riplet, which binds to and adds K63-linked ubiquitin chains to RIG-I, but not MDA5 (18–21). Once activated, RIG-I and MDA5 signal to the adaptor protein MAVS to drive a signal transduction cascade that induces the phosphorylation of IRF3 and then the transcriptional induction of interferon (IFN)-β. HCV infected can also be sensed by TLR3, which signals via TRIF and IRF3 to induce antiviral innate immunity (22). During HCV infection, NS3-NS4A cleaves and/or inactivates MAVS (10, 12, 23, 24), TRIF (25) and Riplet (19) to block IRF3 activation (26).

Here, we aimed to uncouple the roles of NS3-NS4A in replication and immune evasion. We focused on the NS4A transmembrane domain and found a residue, Y16, that regulates differential inactivation of MAVS and Riplet, revealing a new branch of innate immune signaling that controls HCV infection.

## Results

### A Y16F substitution in NS4A disrupts replication of an HCV subgenomic replicon in Huh7 cells, but not in Huh-7.5 cells

The transmembrane domain of NS4A contains two aromatic amino acids: a tryptophan at position 3 (W3) and a tyrosine at position 16 (Y16) (**Fig. 1A**). These two aromatic amino acids, which are conserved in all sequenced HCV strains in the Los Alamos HCV sequence database ((42) and **Fig. 1A**), are located at each end of the NS4A transmembrane domain at the lipid bilayer interface (5, 7). Interestingly, aromatic residues at the termini of transmembrane domains are often important for positioning membrane proteins within lipid bilayers (43–45). Therefore, we hypothesized that these residues may play a role in the proper localization and/or function of the NS3-NS4A protease complex during HCV infection. While both the W3 and the Y16 residues in NS4A are conserved across the eight known HCV genotypes, we chose to focus specifically on the Y16 residue (**Fig. 1A**), with the goal of uncoupling the function of Y16 in HCV replication from targeting of innate immune substrates, such as MAVS and Riplet. As a prior study found that a Y16A substitution inhibited HCV replication (7), we made the more conservative phenylalanine mutation (Y16F) to maintain aromaticity at this position. Here, we analyzed the role of this amino acid in regulating HCV replication and innate immune regulation by NS3-NS4A.

**Figure 1.**
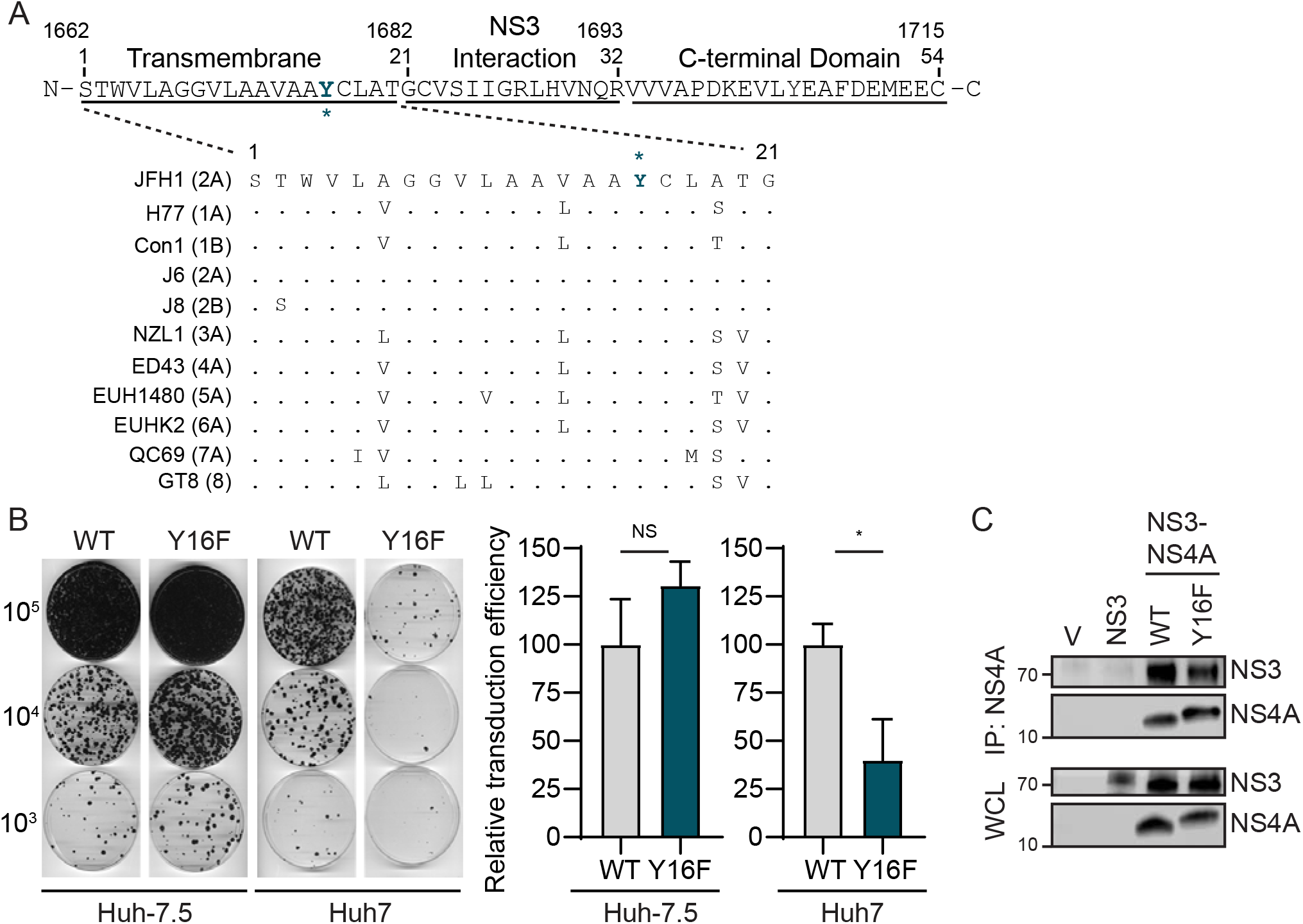
A Y16F substitution in NS4A disrupts replication of an HCV subgenomic replicon in Huh7 cells, but not in Huh-7.5 cells. **(A)** Amino acid sequence of NS4A, with the Y16 residue starred and indicated in teal. Numbers correspond with the amino acid position within NS4A (aa 1-54) or the full-length HCV polyprotein (aa 1662-1715). Strain names are listed as found in the Los Alamos HCV sequence database. Conserved amino acids are indicated with a dot, while differences are listed. **(B)** Representative images of Huh-7.5 or Huh7 cells electroporated with *in vitro* transcribed HCV subgenomic replicon RNA (HP, genotype 1B; WT or Y16F). Cells were plated in serial dilutions (2 × 10^5^, 2 × 10^4^, 2 × 10^3^) and then stained with crystal violet after three weeks of G418 selection. Graphs show the relative transduction efficiency, which denote the % of colonies in Y16F transduced cells relative to WT. Bars indicate mean ± SEM (n = 3-4 biological replicates), with data analyzed by Student’s *t*-test; *p < 0.05, NS = not significant. **(C)** Immunoblot analysis of anti-NS4A immunoprecipitated extracts or whole cell lysate (WCL) from 293T cells transfected with the indicated HCV proteins (genotype 1B) or vector (V). Panels are representative of three independent experiments.

To determine if the Y16F substitution in NS4A altered HCV replication, we first engineered this amino acid change into an HCV replicon encoding a G418 marker (HCV genotype 1B subgenomic replicon; HP replicon (15)). Following *in vitro* transcription, wild-type (WT) or Y16F HCV replicon RNA was electroporated into either liver hepatoma Huh-7.5 cells, which do not have functional RIG-I signaling due to the T55I mutation (15), or Huh7 cells, which have functional RIG-I signaling. In the Huh-7.5 cells, the number of G418-resistant colonies in the WT versus the Y16F HCV replicon-transduced cells was equivalent, indicating that WT and Y16F replicated similarly. However, in Huh7 cells, the Y16F HCV replicon had a reduced transduction efficiency (~3-fold) compared to the WT HCV replicon (**Fig. 1B**). As control, we also measured the interaction of NS4A WT or Y16F with NS3 by co-immunoprecipitation and found that the Y16F substitution did not alter the interaction of NS4A with NS3, nor the ability of the NS3-NS4A protease to process the NS3-NS4A polyprotein junction (**Fig. 1C**). Together, these data reveal that the Y16F mutation results in reduced HCV replication in Huh7 cells, but not Huh-7.5 cells, suggesting that NS4A Y16F may regulate RIG-I-mediated innate immune signaling to promote HCV immune evasion and replication.

### RIG-I deletion in Huh7 cells does not restore HCV NS4A Y16F viral replication

To determine if the Y16F substitution in NS4A specifically altered HCV replication in Huh7 cells during infection, we engineered the NS4A Y16F substitution into the full-length HCV infectious clone (JFH1, genotype 2A (33)). We generated low-passage viral stocks and confirmed that the Y16F mutation was maintained in the resulting virus by PCR amplification of the NS4A region and Sanger sequencing. We then infected Huh-7.5 or Huh7 cells with the HCV WT or Y16F virus, harvested protein lysates over a time course of infection, and measured HCV NS5A protein expression by immunoblot. We found that HCV NS5A protein levels were equivalent in Huh-7.5 cells infected with WT or Y16F HCV (**Fig. 2A**). However, in Huh7 cells, the level of NS5A protein from the Y16F virus was reduced as compared to WT HCV (**Fig. 2B**). In addition to RIG-I, there are likely other genetic differences between Huh7 and Huh-7.5 cells. Thus, to determine if RIG-I was the factor accounting for the differential replication observed between WT and Y16F HCV in Huh7 cells versus Huh-7.5 cells, we generated Huh7-RIG-I knockout (KO) cells using CRISPR/Cas9 genome editing. These Huh7-RIG-I KO cells contain a 252 nucleotide deletion that removes the start codon, preventing RIG-I protein expression (**Fig. 2C**). To confirm a loss of RIG-I signaling, we infected Huh7-RIG-I KO cells with Sendai virus (SV), a virus known to activate RIG-I signaling(15, 46), and observed no SV-mediated induction of RIG-I protein or signaling to the IFN-β promoter, which was restored upon over-expression of RIG-I (15, 16) (**Fig. 2D**).

**Figure 2.**
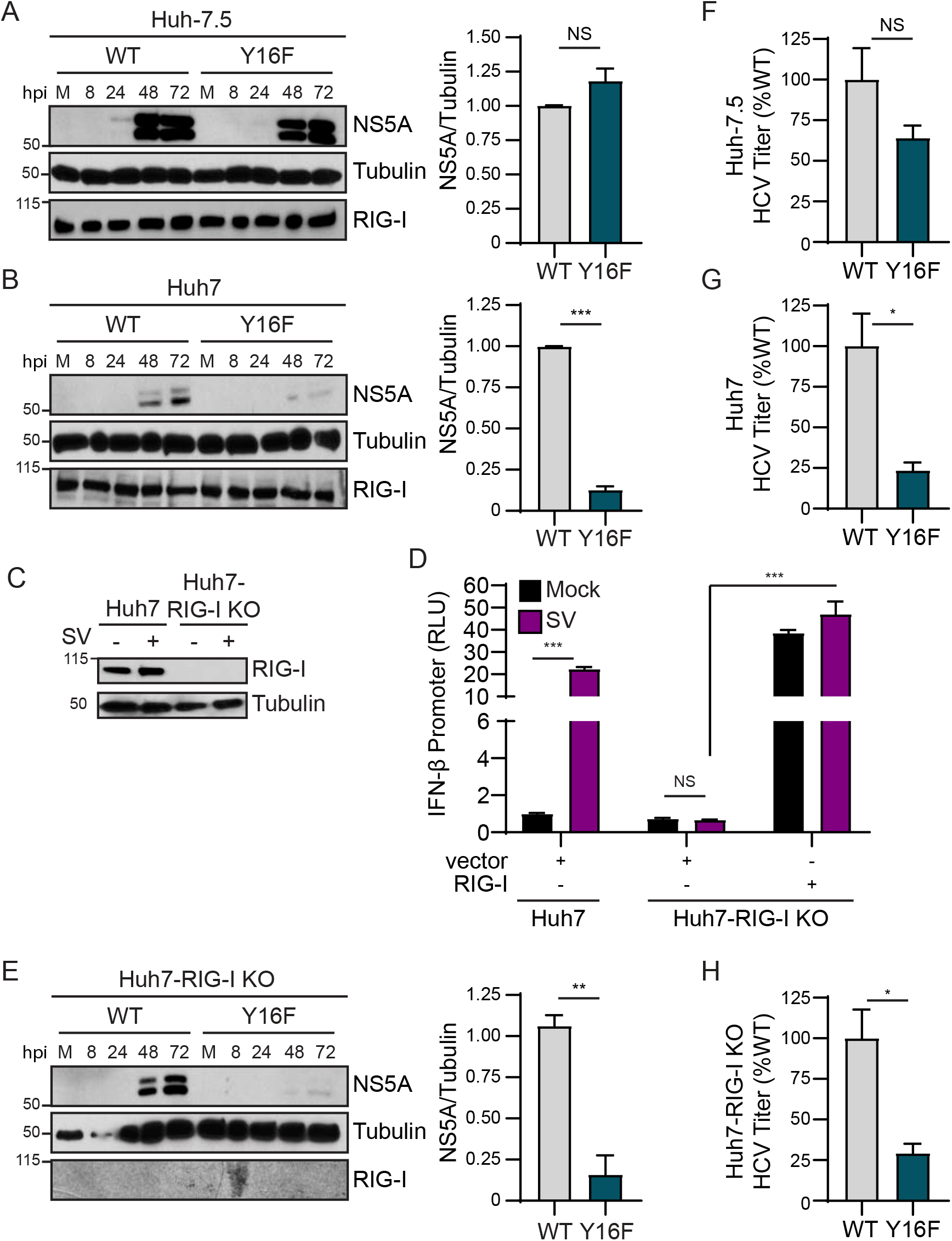
RIG-I deletion in Huh7 cells does not restore HCV NS4A Y16F replication. Huh-7.5 cells **(A)** or Huh7 cells **(B)** were infected with HCV, WT or NS4A Y16F (JFH1, MOI 0.3). Immunoblot analysis was performed on lysates extracted at the indicated hours post infection (hpi) or mock (M). Graphs next to each blot (here, and in **(E)**) show quantification of NS5A protein relative to Tubulin at 72 hpi (mean ± SEM; n = 3 biological replicates). **(C)** Immunoblot of extracts of Huh7 and Huh7-RIG-I KO cells that were mock- or Sendai virus (SV)-infected (20 h). **(D)** IFN-β promoter reporter luciferase expression of Huh7 and Huh7-RIG-I KO cells expressing either vector or full-length RIG-I that were either mock- or SV-infected (20 h). Values show the mean ± SD (n = 3 technical replicates) in relative luciferase units (RLU). **(E)** Huh7-RIG-I KO cells were infected with HCV, WT or NS4A Y16F (JFH1, MOI 0.3). Immunoblot analysis was performed on lysates extracted at the indicated times or mock (M). **(F-H)** Focus forming assay of supernatants harvested from Huh-7.5 **(F)**, Huh7 **(G)**, and Huh7-RIG-I KO **(H)** cells at 72 hpi (MOI 0.3). Data are presented as the percent HCV titer from Y16F relative to the WT (set at 100%) and show the mean ± SEM (n = 3 biological replicates). Data were analyzed by Student’s *t*-test; *p < 0.05, **p < 0.01, ***p < 0.005, NS = not significant.

We next infected these Huh7-RIG-I KO cells with either WT or Y16F HCV and measured HCV NS5A expression from lysates harvested over a time course of infection by immunoblotting. Surprisingly, we found that NS5A protein level from Y16F HCV was not restored to the level of WT in the Huh7-RIG-I KO cells (**Fig. 2E**). We then compared the production of infectious virus from the WT and Y16F viruses in each of these cell lines. In these assays, the supernatants of infected cells were used to infect naïve Huh-7.5 cells to determine the viral titer, which ultimately measures a second round of infection. We found that the while the Y16F virus harvested from Huh-7.5 cells resulted in a somewhat lower level of infectious virus as compared to WT (~40% lower), its level of infectious virus harvested from Huh7 or Huh7-RIG-I KO cells was significantly lower as compared to WT (now ~75% lower) (**Figs. 2F-2H**). Taken together, these data suggest that NS4A Y16 regulates a RIG-I-independent signaling pathway that is non-functional in Huh-7.5 cells.

### HCV NS3-NS4A Y16F retains the ability to cleave MAVS

As NS4A Y16 is located at the membrane lipid bilayer interface (5, 7), and NS4A membrane interactions regulate the molecular mechanisms by which the NS3-NS4A protease targets substrates (7), we hypothesized that the Y16F substitution in NS4A may regulate NS3-NS4A cleavage of MAVS. To test this, we co-expressed NS3-NS4A with Flag-tagged MAVS and found that both NS3-NS4A WT and Y16F cleaved MAVS, while NS3-NS4A containing a mutation that inactivates the protease active site (S139A; SA) did not (**Fig. 3A**). We also found that MAVS cleavage was similar following HCV WT and Y16F infection in both Huh-7.5 and Huh7 cells (**Fig. 3B**). Together, this reveals that the NS4A Y16F substitution does not alter MAVS cleavage by NS3-NS4A.

**Figure 3.**
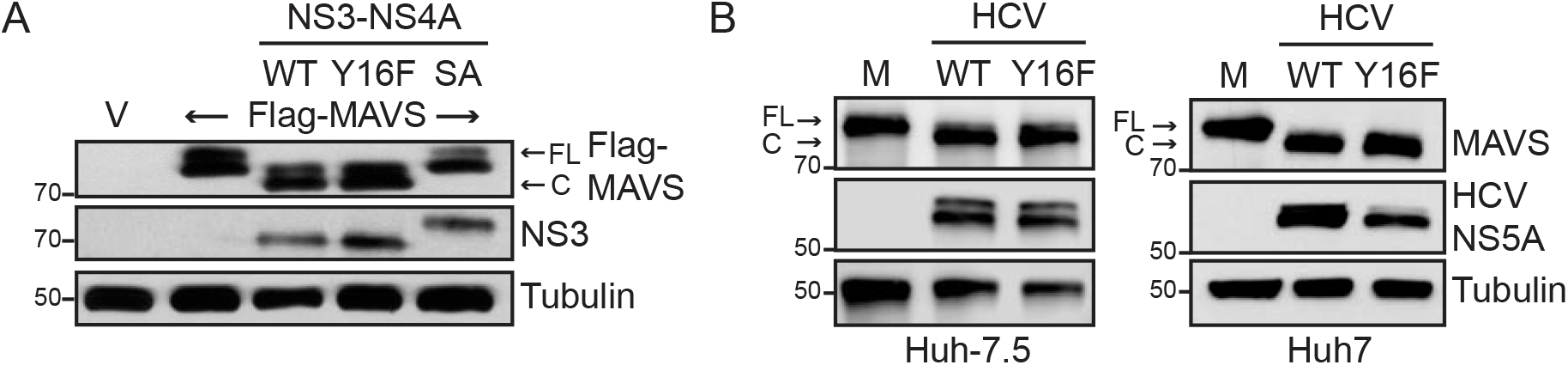
HCV NS3-NS4A Y16F retains the ability to cleave MAVS. **(A)** Immunoblot analysis of lysates harvested from Huh7 cells expressing NS3-NS4A (WT, Y16F, or SA (NS3 active site mutant S139A)) and vector (V) or Flag-MAVS. Arrows indicate the full-length (FL) and cleaved (C) forms of MAVS. **(B)** Immunoblot analysis of lysates harvested at 72 hpi from Huh-7.5 or Huh7 cells that were either mock-infected (M) or infected with HCV, WT or NS4A Y16F (JFH1, MOI 0.3). Arrows indicate the full-length (FL) and cleaved (C) forms of MAVS. Immunoblots are representative of three independent experiments.

### IRF3 deletion in Huh7 cells restores HCV Y16F replication to the levels of HCV WT

We next wanted to determine if the signaling pathway that inhibits HCV Y16F replication requires the IFN-β transcription factor IRF3 (reviewed in (47)). We first generated Huh7-IRF3 KO cells using CRISPR/Cas9 genome editing and determined IRF3 expression and function in these cells by sequencing the IRF3 genetic locus, analyzing IRF3 expression by immunoblot, and confirming that loss of IRF3 prevented SV-mediated antiviral signaling to the IFN-β promoter and that this signaling was restored by IRF3 over-expression (**Figs. 4A-4B**). To determine if IRF3 regulates HCV Y16F replication, we infected Huh7 or Huh7-IRF3 KO cells with either HCV WT or Y16F, measured HCV NS5A expression by immunoblot, and measured release of infectious virus by focus forming assay. While the levels of NS5A expression and infectious Y16F virus were reduced relative to the WT in parental Huh7 cells, as before, these levels were restored to that of WT virus in Huh7-IRF3 KO cells (**Figs. 4C-4D**). Together, these data reveal that NS4A Y16 regulates an IRF3-dependent signaling pathway that can inhibit HCV replication.

**Figure 4.**
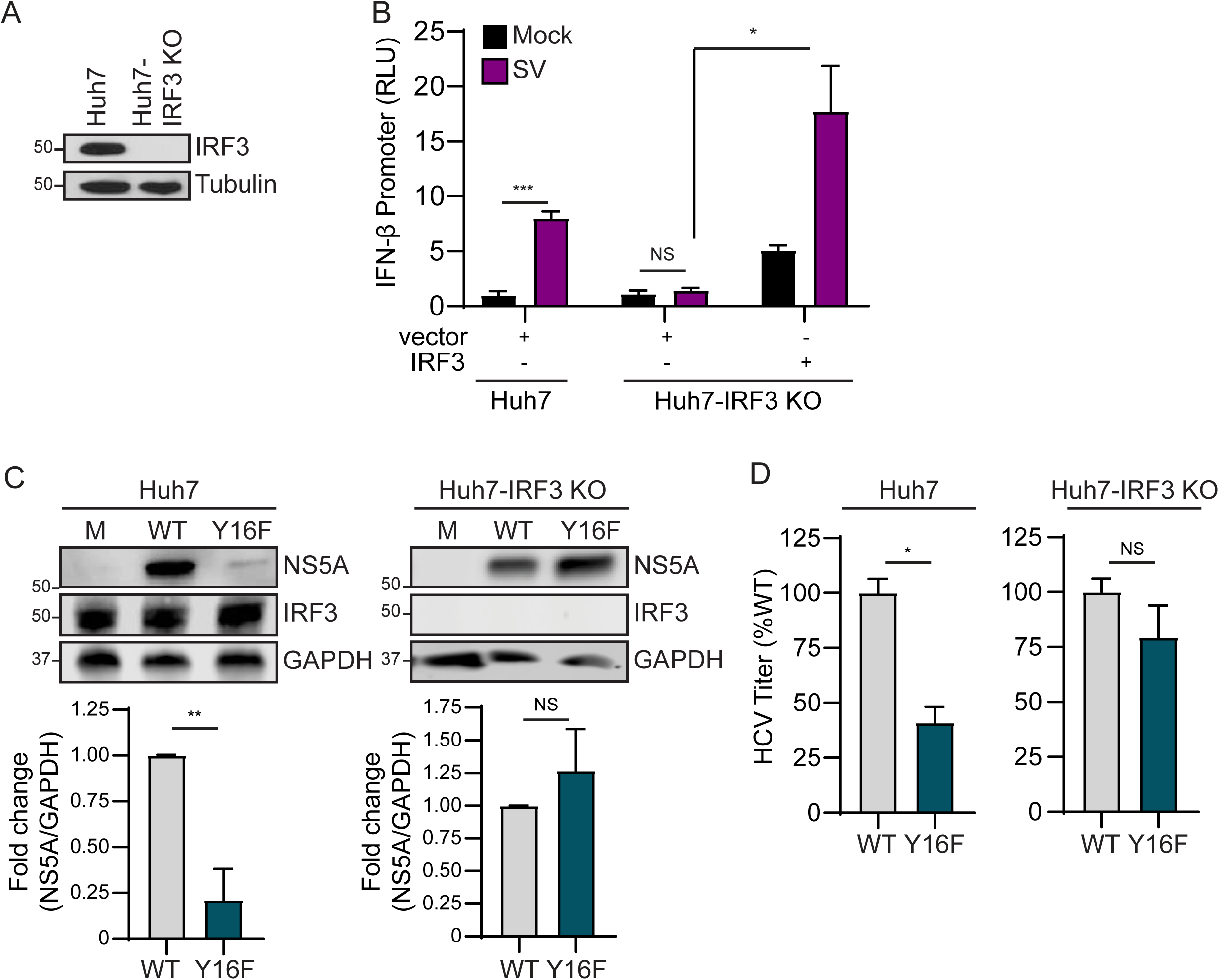
IRF3 deletion in Huh7 cells restores HCV Y16F replication to the levels of HCV WT. **(A)** Immunoblot of extracts of Huh7 and Huh7-IRF3 KO cells. **(B)** IFN-β promoter reporter luciferase expression of Huh7 and Huh7-IRF3 KO cells expressing either vector or full-length IRF3 that were either mock- or SV-infected (20 h). Values show the mean ± SD (n = 3 technical replicates). in relative luciferase units (RLU). **(C)** Immunoblot analysis of lysates harvested at 72 hpi from Huh7 and Huh7-IRF3 KO cells infected with HCV, WT or NS4A Y16F (JFH1, MOI 0.3). Graphs below each blot show quantification of NS5A protein relative to GAPDH (mean ± SEM; n = 3 biological replicates). **(D)** Focus forming assay of supernatants harvested at 72 hpi from Huh7 or Huh7-IRF3 KO cells infected with HCV, WT or NS4A Y16F (MOI 0.3). Data are presented as the percent of HCV titer from Y16F relative to WT (set at 100%) and show the mean ± SEM (n = 2 biological replicates). Data were analyzed by Student’s *t*-test; *p < 0.05, **p < 0.01, ***p < 0.005, NS = not significant.

### HCV NS3-NS4A Y16F does not block IRF3 activation

As our data suggested that HCV Y16F replication was inhibited by IRF3-mediated signaling, we hypothesized that NS3-NS4A Y16F would be unable to block IRF3 activation. During viral infection, IRF3 is activated by a multi-step process, including phosphorylation by the kinases TBK1 and IKKε, resulting in dimerization, and finally translocation from the cytosol to the nucleus, where it activates transcription of IFN-β (41). Importantly, it is well-known that over-expression of the WT NS3-NS4A protease can block this nuclear translocation of IRF3 in response to virus infection (12, 48). Therefore, we measured the ability of WT or Y16F NS3-NS4A to block the nuclear translocation of GFP-IRF3 in response to SV. GFP-IRF3 translocated to the nucleus in approximately 50% of the SV-infected cells, as measured by immunofluorescence assay (**Figs. 5A-5B**). While the NS3-NS4A WT blocked nearly all of this nuclear translocation, NS3-NS4A Y16F did not (**Figs. 5A-5B**), revealing that NS3-NS4A Y16F has a reduced ability to inhibit IRF3 activation.

**Figure 5.**
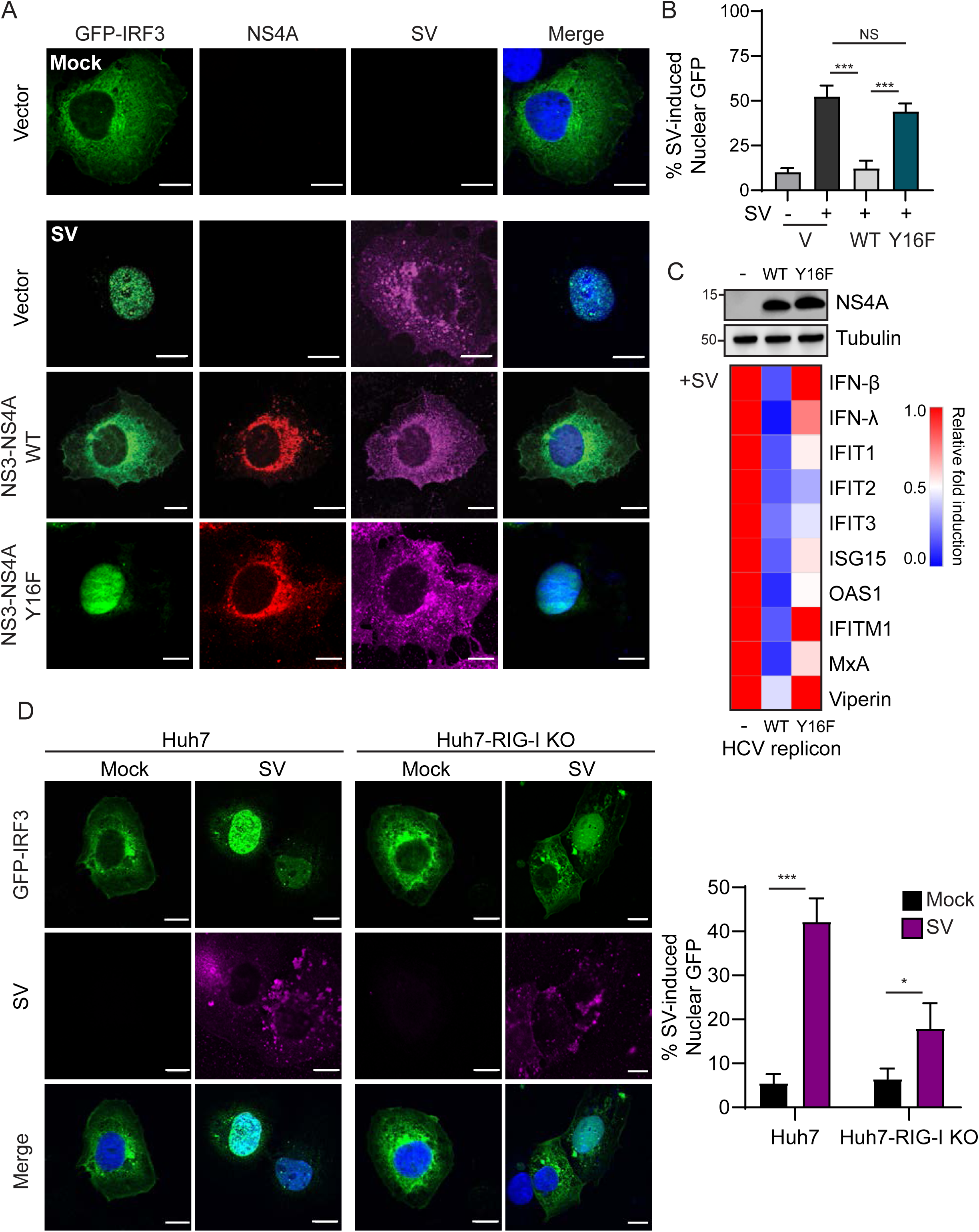
HCV NS3-NS4A Y16F does not block IRF3 activation. **(A)** Confocal micrographs of Huh7 cells expressing GFP-IRF3 (green) and either NS3-NS4A WT or Y16F (genotype 1B), or vector, that were either mock- or SV-infected (20 h) and immunostained with anti-NS4A (red) or anti-SV (magenta). Nuclei were stained with Hoescht (blue). Scale bar: 10 μm. **(B)** Quantification of the percent of cells both expressing GFP-IRF3 and positive for SV. Data are displayed as mean ± SEM (n = three biological replicates of 50-100 cells counted in each condition and replicate). Data were analyzed by one-way ANOVA; ***p < 0.005. **(C)** Immunoblot analysis of lysates from Huh7 (-), Huh7-HP WT replicon, or Huh7-HP Y16F replicon cells, and a heatmap (below) that shows the mean relative fold induction (SV-infected/mock-infected, relative to *HPRT1*) of specific genes as measured by RT-qPCR analysis of RNA from mock- or SV-infected (20 h) Huh7, Huh7-HP WT replicon, or Huh7-HP Y16F replicon cells from three biological replicates. (*D*) Confocal micrographs of Huh7 and Huh7-RIG-I KO cells expressing GFP-IRF3 (green) that were either mock- or SV-infected (20 h) and immunostained with anti-SV (magenta). Nuclei were stained with Hoescht (blue). Scale bar: 10 μm. Graph shows the quantification of the percent of cells both expressing GFP-IRF3 and positive for SV. Data are displayed as mean ± SEM (n = three biological replicates of 50-100 cells counted in each condition and replicate) and were analyzed by Student’s *t*-test; *p < 0.05 and ***p < 0.005.

To test if NS3-NS4A Y16F similarly did not block IRF3 activation in the context of HCV replication, we utilized the HCV replicon system, which activates RIG-I signaling but prevents the transduction of IRF3 signaling by NS3-NS4A cleavage of MAVS, to prevent HCV or SV-induced innate immune signaling (48). We infected control cells and cells stably expressing either WT or Y16F subgenomic replicons with SV and then measured induction of IFN-β and several ISGs by RT-qPCR. While the WT HCV replicon prevented SV-mediated induction of all ISGs tested, the HCV Y16F replicon did not block induction of IFN-β, IFN-λ, IFITM1, and Viperin, and only partially blocked induction of several other ISGs (**Fig. 5C**). Interestingly, in the Huh7-RIG-I KO cells, GFP-IRF3 translocated to the nucleus in approximately 15%-20% of SV-infected cells, while only being nuclear in less than 10% of mock-infected cells, suggesting that other signaling molecules are capable of activating IRF3 in the absence of RIG-I (**Fig. 5D**). Taken together, these data reveal that the Y16F substitution prevents NS3-NS4A from fully blocking IRF3 activation and signaling in response to viral infection.

### HCV NS4A Y16F does not target Riplet

Our data described thus far reveal that NS4A Y16 regulates NS3-NS4A inhibition of IRF3-mediated antiviral signaling. This IRF3-mediated signaling, which limits HCV replication, is RIG-I-independent and MAVS-cleavage independent. Together, these data suggest: (1) that there is a factor that induces signaling to IRF3 that is targeted by NS4A Y16 (and not Y16F), and (2) that this factor is present in Huh7 cells but absent or non-functional in Huh-7.5 cells. NS3-NS4A cleaves and inactivates three known host proteins involved in the IRF3 signaling axis: MAVS, TRIF (the TLR3 signaling adaptor), and Riplet (10, 12, 19, 24, 25, 48). Since we have demonstrated that NS3-NS4A Y16F cleaves MAVS (**Fig. 3**), and it is known that Huh7 cells do not have functional TLR3 signaling (49), we hypothesized that the E3 ubiquitin ligase Riplet may be differentially regulated by NS3-NS4A WT and Y16F. Interestingly, we found that Huh-7.5 cells express reduced levels of Riplet (*RNF135*) mRNA as compared to Huh7 cells (**Fig. 6A**). This low level of Riplet likely renders it incapable of driving signaling. Therefore, we tested if Riplet ectopic expression in Huh-7.5 cells could limit HCV Y16F replication relative to HCV WT. We generated Huh-7.5 cells expressing V5-tagged Riplet (**Figs. 6A-6B**), infected these cells with HCV WT or Y16F, and measured HCV NS5A expression. In Huh-7.5 + Riplet-V5 cells, but not Huh-7.5 cells, HCV Y16F replication was reduced compared to WT (**Fig. 6B**). Similarly, the amount of infectious virus generated in the Huh-7.5 + Riplet-V5 cells or Huh7 cells from the HCV Y16F virus was also much lower than WT (~90% lower in each), but in the Huh-7.5 cells, the level of Y16F virus was still only partially reduced compared to WT, similar to before (~50% lower) (**Fig. 6C, Fig. 2**). We note that the overall levels of HCV replication (both WT and Y16F) in the Huh-7.5 + Riplet-V5 cells were lower than those seen in the parental Huh-7.5 cells, likely due to the higher levels of Riplet expression in these cells (**Fig. 6**) and the known role of Riplet in inhibiting HCV replication (19).

**Figure 6.**
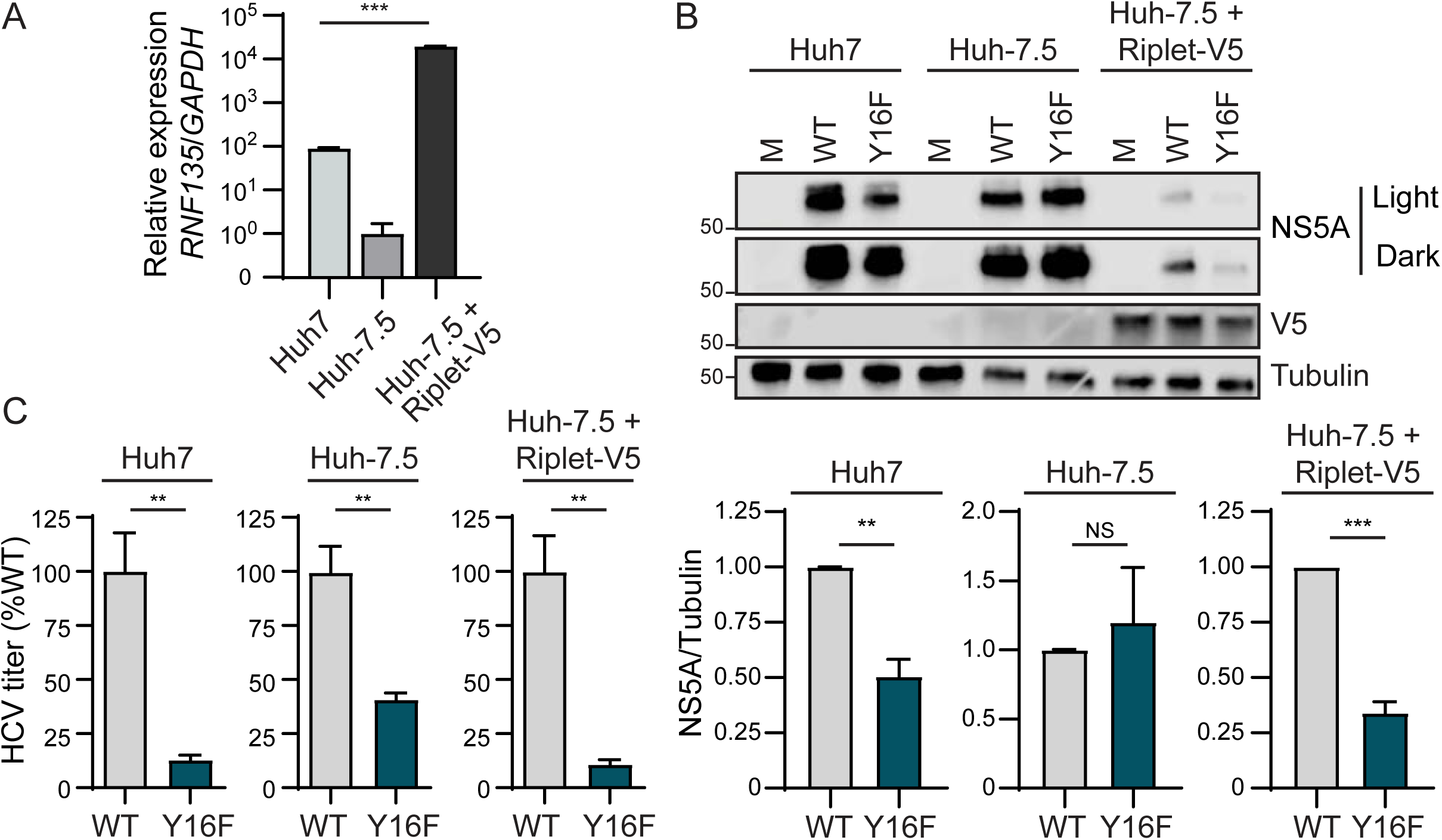
Over-expression of Riplet reduces HCV NS4A Y16F replication in Huh-7.5 cells. **(A)** *RNF135* (Riplet) expression relative to *GAPDH* from Huh7, Huh-7.5, and Huh-7.5 + Riplet-V5 cells, as analyzed by RT-qPCR, with data displayed as mean ± SD (n = 2-3 technical replicates). Data were analyzed by one-way ANOVA analysis across the means of the three groups. **(B)** Immunoblot analysis of lysates harvested from the indicated cell lines infected with HCV, WT or NS4A Y16F (JFH1, MOI 0.3), or mock-infected (M), at 72 hpi. Two different exposures (Light and Dark) are shown for NS5A. Graphs below each blot show mean ± SEM (n= 3 biological replicates) of quantification of NS5A protein relative to Tubulin. **(C)** Focus forming assay of supernatants harvested at 72 hpi from the indicated cell lines infected with HCV, WT or NS4A Y16F (MOI 0.3). Data are presented as the percent HCV titer from Y16F relative to the WT (set at 100%) and show the mean ± SEM (n = 3 biological replicates). Data were analyzed by Student’s *t*-test; *p < 0.05, **p < 0.05, ***p < 0.005, NS = not significant.

To test the role of NS4A Y16 in targeting Riplet, we first examined the localization of over-expressed NS3-NS4A WT or Y16F with HA-tagged Riplet in Huh7 cells by immunofluorescence. Similar to others, we did not detect any major difference in the localization of NS4A WT or Y16F (5). In cells expressing NS3-NS4A WT, we found that Riplet was localized in small, punctate aggregates throughout the cytoplasm, whereas in cells expressing NS3-NS4A Y16F, Riplet was diffusely localized throughout the cytoplasm, similar to that seen in vector-expressing cells and described previously (18) (**Fig. 7A**). We also found that in cells expressing NS3-NS4A WT, but not Y16F, Riplet and NS4A were in close proximity to each other (**Fig. 7A, zoom**), suggesting that NS4A may interact with Riplet in a Y16-dependent manner. Indeed, we found that NS4A alone interacted with Flag-tagged Riplet and that the Y16F mutation reduced this interaction by approximately 70% (**Fig. 7B**). Taken together, these data suggest that the NS4A Y16 residue is necessary for the ability of NS3-NS4A to interact with Riplet and to block antiviral innate immune signaling during HCV infection

**Figure 7.**
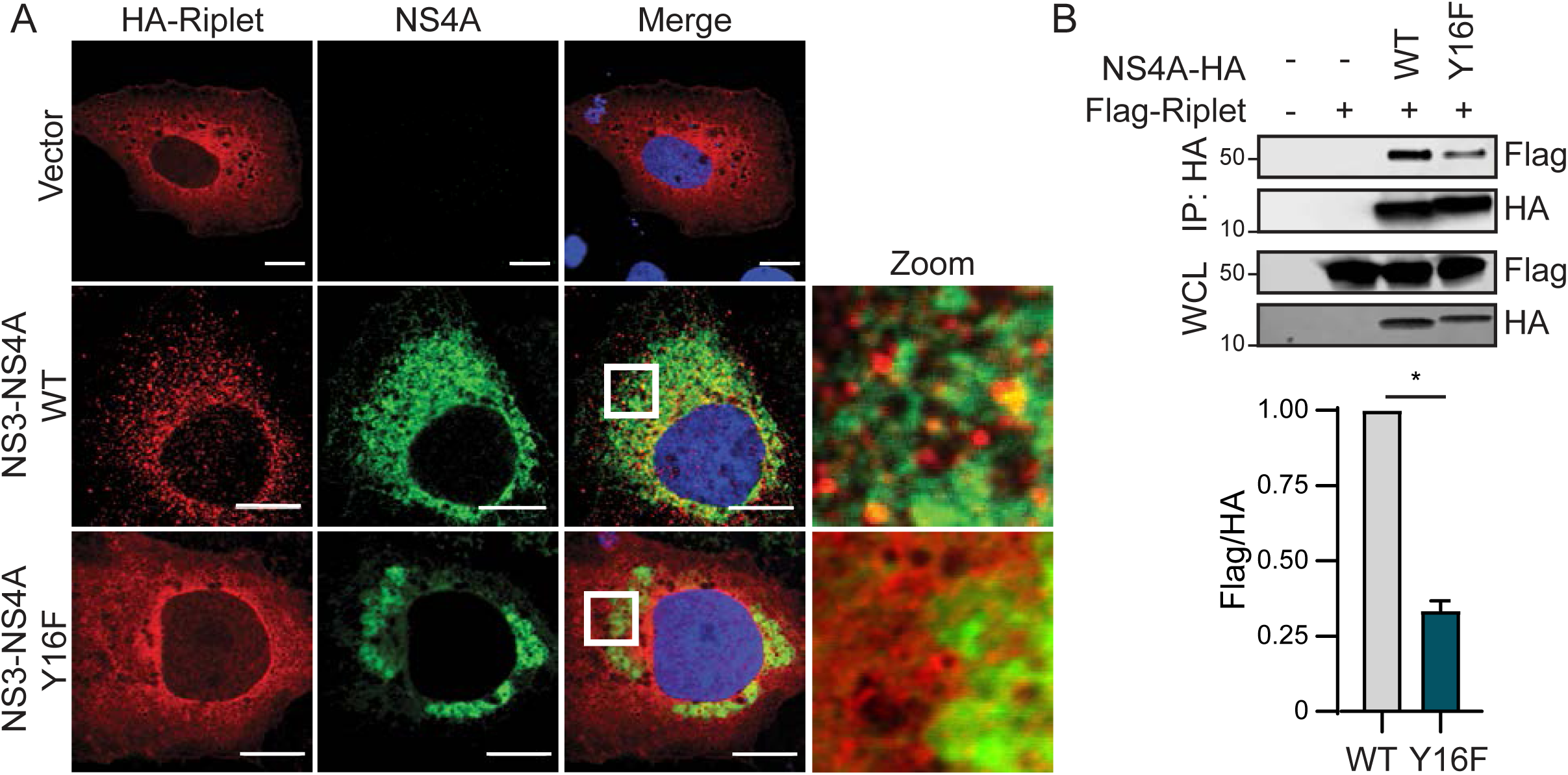
HCV NS4A interaction with Riplet is reduced with Y16F mutation. **(A)** Confocal micrographs of Huh7 cells expressing HA-Riplet and either NS3-NS4A WT or Y16F (genotype 1B), or vector, that were immunostained with anti-NS4A (green) and anti-HA (red), with the nuclei stained with Hoescht (blue). Zoom panel is taken from the images in the white boxes. Images are representative of ~50 cells analyzed. Scale bar: 10 μm. **(B)** Immunoblot analysis of anti-HA (NS4A) immunoprecipitated extracts or whole cell lysate (WCL) from Huh7 cells transfected with plasmids expressing Flag-Riplet and NS4A-HA (genotype 1B) WT or Y16F, or vector (-). The graph directly below shows the mean ± SEM (n = 3 biological replicates) of the relative fold change of Flag-Riplet to NS4A-HA in the immunoprecipitated lanes. Data were analyzed by Student’s *t*-test; *p < 0.05.

## Discussion

Our results identify a new antiviral signaling program regulated by HCV NS3-NS4A. We found that mutation of NS4A Tyr-16 to phenylalanine, in both the context of an HCV subgenomic RNA replicon and in the context of fully infectious HCV, results in reduced viral replication in Huh7 cells, but not in Huh-7.5 cells. We show that both NS3-NS4A WT and Y16F cleave MAVS. Further, we found that Huh-7.5 cells, in addition to lacking RIG-I signaling (15), express low levels of Riplet (**Fig. 6**). Importantly, ectopic expression of Riplet in Huh-7.5 cells resulted in reduced replication of HCV Y16F compared to WT virus. We also found that NS4A WT binds to Riplet, while NS4A Y16F does not bind as well. Taken together, this supports the model that HCV inactivates Riplet to prevent signaling to IRF3 and an antiviral response that can inhibit HCV replication. Our work reveals that the NS3-NS4A Y16 residue plays a critical role in the inactivation of this signaling pathway. Thus, NS4A Y16 regulates an antiviral signaling program activated by a Riplet-IRF3-dependent, but RIG-I-MAVS-independent, signaling axis.

We found that HCV containing an NS4A Y16F substitution in two HCV genotypes, either the JFH1 genotype 2A virus or the HP genotype 1B subgenomic replicon, has lower levels of replication than WT in Huh7 cells (**Fig. 1B, Fig. 2A and 2B**), but that both the WT and Y16F viruses have similar levels of replication in Huh-7.5 cells or in Huh7-IRF3 KO cells (**Fig. 2B, Figs. 4C-4D**). Although we did find that in experiments that assessed viral titer, the Y16F virus from Huh-7.5 cells had a reduced viral titer as compared to the WT. However, this reduction (~50%) was not as much as that of virus harvested from the Huh7 parental, Huh7-RIG-I KO, or Huh-7.5 + Riplet-V5 cells (~80%). This, along with our replication experiments, suggests that the while the Y16F substitution does not itself directly affect the functions of the HCV protease in replication, including HCV polyprotein processing, NS3 helicase function, or viral assembly and envelopment, the virus with this substitution may still be inhibited by the low, remaining levels of Riplet present in the Huh-7.5 cells that we use to measure the production of infectious virus. Indeed, the viral titers between the WT and Y16F viruses harvested from the Huh7-IRF3 KO cells were similar to each other (**Fig. 4D**). Similar to our findings, Kohlway and colleagues found that the replication of genotype 2A subgenomic replicon (pYSGR-JFH1/GLuc) containing this Y16F substitution was not altered in Huh-7.5 cells (7), while Brass and colleagues did find reduced replication of a Y16F genotype 1B subgenomic replicon (pCon1/SG-Neo(I)/AflII) in Huh-7.5 cells (5). While it is unclear what mediates the difference in our HCV replication results from those of Brass and colleagues, it is possible that this could be due to differences in the replication fitness of the replicons used or that Huh-7.5 cells from different labs do not have the same expression levels of Riplet. Unfortunately, all of our attempts to use CRISPR to delete Riplet from Huh7 cells were unsuccessful. Nevertheless, as we found that the Y16F substitution does not affect either the interaction of NS4A with NS3, processing of the NS3/NS4A junction, or MAVS cleavage, our results suggest that it has a specific role in targeting NS3-NS4A to Riplet.

The mechanisms by which the HCV protease targets and inactivates Riplet are not entirely clear. Riplet is an E3 ubiquitin ligase localized in the cytoplasm that activates RIG-I by both binding and adding K63-linked ubiquitin chains to it (20, 50). While others have concluded that NS3-NS4A cleaves Riplet in the first amino acid of its RING domain resulting in its destabilization (19), we were not able to detect a Riplet cleavage product or a reduction in Riplet protein abundance by immunoblot analysis upon over-expression of NS3-NS4A in cells, although we cannot rule out this possibility. While it is possible that NS3-NS4A inactivation of Riplet via cleavage may result in its destabilization, analogous to how NS3-NS4A cleavage of TRIF accelerates its proteolysis (25), it is also possible that simply the binding of NS3-NS4A to Riplet can inactivate it. Indeed, we did find that the localization of Riplet changed from cytoplasmic to punctate, often near NS4A, following over-expression of NS3-NS4A WT, but not Y16F, which could either represent a differential localization as a result of binding to NS4A to prevent Riplet function or represent cleavage by the WT NS3-NS4A (**Fig. 7A**). Indeed, the dengue virus protease co-factor NS2 (analogous to HCV NS4A) inactivates cGAS simply by binding to it and inducing its autophagic degradation (51). Additionally, the influenza A virus NS1 protein inactivates Riplet by binding to it (37). Therefore, while it is clear that NS3-NS4A inactivates Riplet, further studies are needed to determine the exact mechanisms by which this occurs.

While HCV NS4A anchors the NS3-NS4A protease to intracellular membranes (6), the mechanisms by which the Y16F substitution in NS4A would specifically alter Riplet localization and block Riplet signaling are unclear. Similar to others, we did not find that the Y16F substitution altered the localization of NS4A within membranes (5). Since NS4A can bind Riplet in the absence of NS3, it is possible that NS4A Y16 is simply required for Riplet binding, either directly or through other proteins. In fact, as the hydroxyl group of this tyrosine residue in NS4A is positioned such that it interacts with the phospholipid head groups of the membrane bilayer, while a phenylalanine at the position would be missing this hydroxyl group, Y16 may be poised to mediate protein-protein interaction directly with Riplet or with accessory binding proteins (5). We also note that it is possible that phosphorylation of NS4A Y16 could regulate these protein-protein interactions. Thus, NS4A Y16 likely mediates interactions with Riplet to prevent Riplet from interacting with proteins that mediate antiviral innate immune signaling.

Our results suggest that HCV activates a Riplet-dependent signaling cascade to IRF3 that is independent of both RIG-I and MAVS. The following pieces of evidence presented within this manuscript support the existence of this pathway: (1) NS3-NS4A WT and Y16F both cleave MAVS (**Fig. 3**), (2) Y16F cannot bind to Riplet as well as WT (**Fig. 7**); (3) NS3-NS4A WT, but not Y16F, blocks SV-mediated IRF3 activation and induction of ISGs (**Fig. 5**); (4) WT and Y16F viruses only grow equivalently to each other in cells that lack both RIG-I and Riplet or lack IRF3 (**Fig. 1-2; Fig. 4**); (5) over-expression of Riplet in cells without RIG-I signaling can reduce Y16F viral replication (**Fig. 6**). While the identification of this RIG-I-MAVS independent signaling cascade that induces IRF3 activation and IFN-β was surprising to us, others have shown that infection of RIG-I KO mouse embryonic fibroblasts with vesicular stomatitis virus, known to be sensed by only RIG-I (52), does result in a small induction of IFN-β mRNA, even though other stimuli do not induce IFN-β in these cells (20). Thus, it is possible that in our human Huh7-RIG-I KO cells, ISGs are induced during HCV infection to limit Y16F viral replication. Indeed, we do see a low level of IRF3 nuclear translocation in response to SV in these cells (**Fig. 5D**). Overall, this induction of this Riplet-IRF3 signaling pathway in the absence of RIG-I is likely stimulus-dependent and cell type-dependent.

We do not yet know the full identity of this Riplet-IRF3 signaling cascade regulated by NS3-NS4A Y16. We predict that Riplet is either directly adding K63-linked ubiquitin chains to signaling proteins in this pathway or that it interacts with these signaling proteins to activate them, as it does with RIG-I (18, 20). The only known Riplet-interacting protein that is K63-ubiquitinated is RIG-I. Therefore, the Riplet-signaling target is likely not MDA5, because it is not a Riplet substrate (18, 20) and the Y16F protease is capable of cleaving the downstream signaling protein. It could be the IRF3 kinases TBK1 and IKKε or some other unknown upstream factor (53, 54). Future studies are needed to further identify the Riplet-interacting proteins that activate this non-canonical antiviral signaling pathway.

Here, we identify a new, non-canonical branch of an antiviral signaling pathway regulated by NS3-NS4A that can inhibit HCV infection. This signaling pathway is driven by Riplet to induce IRF3 activation, and our data suggest that it does not require MAVS. This signaling results in the transcriptional induction of IRF3-regulated genes, including IFN-β and several ISGs. Ultimately, full characterization of this novel pathway may reveal insights into antiviral innate immunity to other RNA viruses, such influenza A virus, which inactivates Riplet (37). In summary, our work identified a specific amino acid in NS4A which uncouples Riplet inactivation from MAVS cleavage and HCV replication. Identification of this residue allowed us to show that Riplet can be regulated independently from RIG-I and MAVS during HCV infection, and that NS3-NS4A regulates a Riplet/IRF3-dependent, RIG-I-MAVS-independent branch of an antiviral signaling pathway that limits HCV infection.

## Materials and Methods

### Cell culture

Huh7 and Huh-7.5 (15) cells (gift of Dr. Michael Gale Jr., University of Washington), as well as 293T cells (ATCC; CRL-3216), were grown at 37°C with 5% CO_2_ in Dulbecco’s modification of eagle’s medium (DMEM; Mediatech) supplemented with 10% fetal bovine serum (FBS; HyClone), and 25 mM 4-(2-hydroxyethyl)-1-piperazineethanesulfonic acid (HEPES; Thermo Fisher), referred to as complete DMEM (cDMEM). Huh7 and Huh-7.5 cells were verified using the Promega GenePrint STR kit (DNA Analysis Facility, Duke University), and all cells were tested and found to be *Mycoplasma-free* using the LookOut PCR Detection Kit (Sigma).

### Plasmids and transfections

These plasmids were used in this study: pEF-NS3, pEF-NS3-NS4A (genotype 1B), pEF-NS3-NS4A S1165A (27); pHCV-HP WT (containing the following 7 amino acid changes: NS3 (P1115L, K1609E), NS4B (Q1737R), NS5A (P2007A, L2198S, S2236P), and NS5B (V2971A)) and pHCV-HP ΔNS5B (28); pEF-Tak-Flag MAVS (12); pCR-BluntII-TOPO (Addgene #41824) (29), phCas9 (Addgene #41815) (29); pCMV-Renilla and pGL4.74 [hRluc/TK] (Promega); pIFN-β-Luc (30); pEF-Tak-Flag and pEF-Tak Flag RIG-I (31); pEGFP-C1-IRF3 (32); psJFHI-M9 WT (33); pX330 (Addgene #42230) (34); pcDNA-Blast (35); pPAX2 and pMD2.G (Addgene #35002 (36); Addgene #12259); pCCSB-Riplet-V5 (Dharmacon: NM_032322.4, cDNA clone MGC161700); pCAGGS-HA-Riplet (37) (Dr. Michaela Gack at the University of Chicago); and pCMV-Flag-IRF3 WT (38). psJFHI-M9 Y16F, pEF-NS3-NS4A Y16F, pHCV-HP-Y16F, were generated by QuikChange site-directed mutagenesis (Stratagene) of psJFHI-M9, pEF-NS3-NS4A, and pHCV-HP. The oligonucleotide sequences used for cloning are listed in Table 1. pEF-Tak Flag-Riplet was generated by InFusion (ClonTech) cloning of pCAGGS-HA-Riplet into pEF-Tak. To generate the RIG-I CRISPR guide RNA plasmids (pCR-BluntII-Topo-sgRIGI-I and pCR-BluntII-Topo-sgRIGI-2), sgRNA oligonucleotides were annealed and inserted into *AflII*-digested pCR-BluntII-Topo by Gibson Assembly (New England Biolabs). To generate the IRF3 CRISPR guide RNA plasmids, sgRNA oligonucleotides were annealed and inserted into *BbsI*-digested pX330. All oligonucleotide sequences are listed in Table 1. The sequences of all plasmids were verified by DNA sequencing and are available upon request. DNA transfections were done using FuGENE6 (Promega) according to manufacturer’s instructions.

**Table 1:**
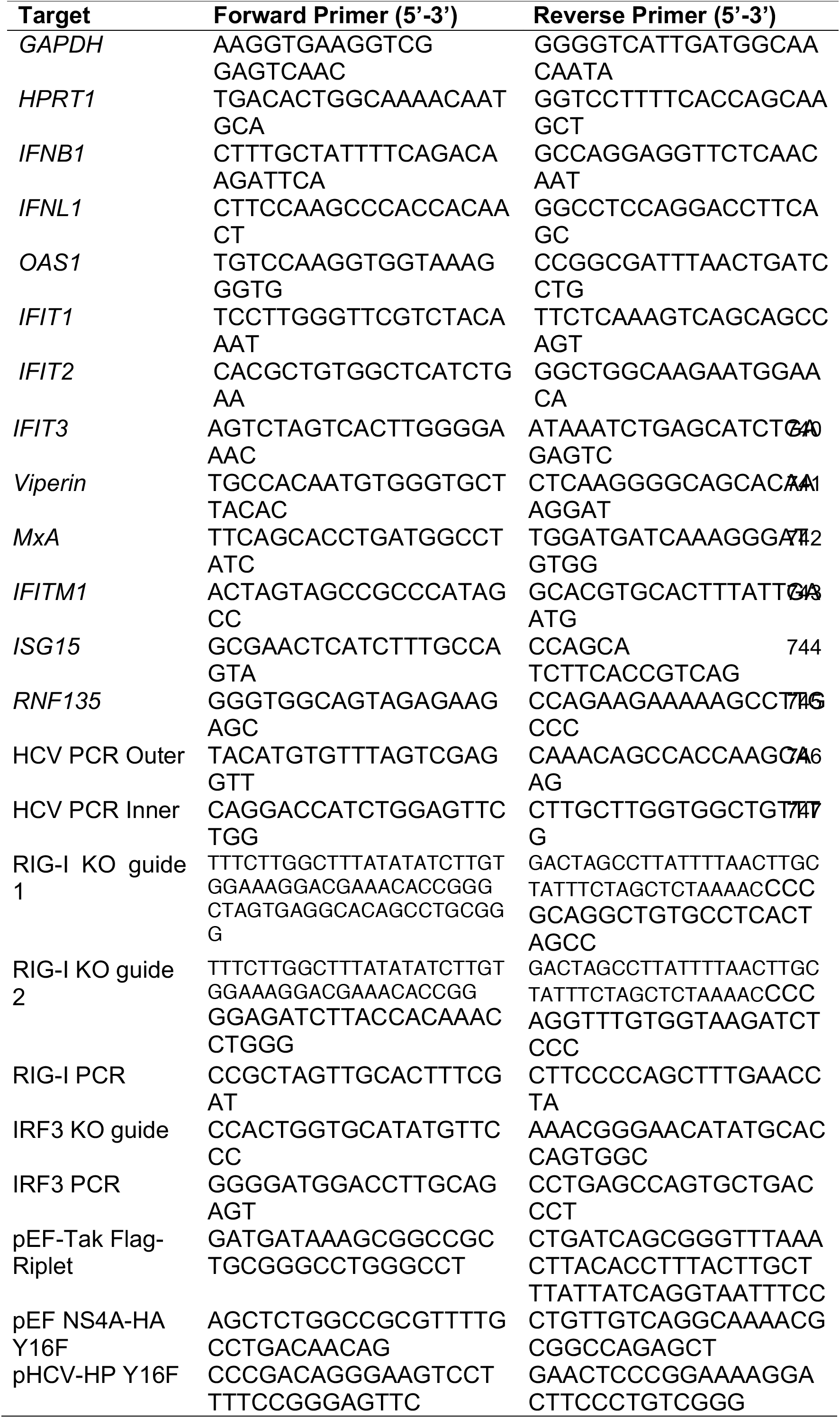
Oligonucleotides used for RT-qPCR and cloning.

### Generation of knock out (KO) cell lines

Huh7-RIG-I KO cells were generated by CRISPR/Cas9, using two guides targeting the intron before exon 1 (sgRNA 1) and within exon 1 (sgRNA 2) that were designed with the CRISPR design tool (http://crispr.mit.edu). pCR-BluntII-Topo-sgRIGI-I and pCR-BluntII-Topo-sgRIGI-2, along with phCas9, which expresses Cas9 and neomycin (G148) resistance, were transfected into Huh7 cells. Huh7-IRF3 KO cells were generated by CRISPR/Cas9 using a single guide that targets exon 2 (39). pX330-sgIRF3, along with pcDNA-Blast (which encodes blasticidin resistance), were transfected into Huh7 cells. In both cases, cells were re-plated the day after transfection at limiting dilutions into 15 cm dishes and then incubated with cDMEM containing either 0.4 mg/ml geneticin (G418; Life Technologies) for 5 days or 0.2 μg/ml blasticidin for 3 days. Individual cell clones were then selected and expanded. Isolated clones were screened for either RIG-I or IRF3 protein expression by immunoblot. Genomic DNA was isolated from candidate RIG-I or IRF3 KO cell clones using the QuickExtract DNA extraction solution (Epicentre). Genomic DNA isolated from the RIG-I or IRF3 KO cell clones was then amplified by PCR using primers spanning exon 1 for RIG-I or exon 2 for IRF3 (see Table 1). The resulting amplicons were cloned into pCR4-TOPO TA (Invitrogen) and Sanger sequenced. For RIG-I, all five of the sequenced genomic DNA clones had the start codon and exon 1 removed (four clones: 252 bp deletion and 1 clone: 250 bp deletion). For IRF3, all five of the clones sequenced had a 4 bp deletion at the beginning of exon 2 that causes a frame shift resulting in a premature stop codon within exon 2.

### Generation of Huh-7.5 + Riplet-V5 cells

To generate Riplet-V5 expressing lentivirus, 293T cells were transfected with pCCSB-Riplet-V5, psPAX2, and pMD2.G. Supernatant was harvested at 48 hours post-transfection and filtered through a 0.45 μm filter. Huh-7.5 cells were then infected with the Riplet-V5 lentivirus (500 μl per well of a 6-well plate), and the next day virus was removed and replaced with cDMEM with 0.2 μg/ml blasticidin until mock-transduced cells died (3-4 days). Blasticidin-resistant cells were harvested as pools, and cells were verified as transduced by immunoblot for Riplet-V5 and RT-qPCR analysis for *RNF135* (Riplet).

### HCV replicons

RNA was *in vitro* transcribed (MEGAscript T7 transcription kit; Thermo Fisher) from ScaI-digested HP-HCV replicon plasmid DNA, either WT, Y16F, or ΔNS5B. The *in vitro* transcribed RNA was treated with DNase (Thermo Fisher), purified by phenol-chloroform extraction, and integrity verified on a denaturing gel. For electroporation, 1 μg of HCV replicon RNA was mixed with 4 × 10^6^ Huh7 or Huh-7.5 cells in cold 1X phosphate buffered saline (PBS) in a 4 mm cuvette and then electroporated at 960 μF and 250 V with a Gene Pulser Xcell system (Bio-Rad). Electroporated cells were plated into 10 cm plates at 2 × 10^5^, 2 × 10^4^, 2 × 10^3^ cells per dish, along with 2 × 10^5^ cells that had been electroporated with ΔNS5B RNA. Four hours post electroporation, cells were washed three times with 1X PBS and then once with cDMEM. At twenty-four hours post electroporation, media was changed to cDMEM supplemented with 0.4 mg/ml G418. Following three weeks of G418 selection, cells were fixed and stained with crystal violet in 20% methanol. Colonies from triplicate plates were counted to determine the relative transduction efficiency, expressed as the percentage of Y16F colonies that were stably transduced relative to WT. Huh7-HP WT and Huh7-HP Y16F replicon cell lines were generated by isolating and expanding single clones. The presence of the HCV replicon was determined by sequencing the NS4A-containing region following cDNA synthesis on extracted RNA (RNeasy RNA extraction kit, Qiagen) and PCR amplification of the NS4A region. Oligonucleotides used for PCR and sequencing are listed in Table 1.

### HCV stock generation and infections

HCV JFH1-M9 WT and Y16F virus stocks were generated as described previously (33). The sequence of the virus at NS4A was confirmed after each passage by sequencing nested PCR products from generated cDNA using the oligonucleotides indicated in Table 1, as previously described (9). For HCV infections, cells were incubated in a low volume of serum-free DMEM containing virus at a multiplicity of infection (MOI) of 0.3 for 2-3 hours, after which cDMEM was replenished. To quantify virus, cellular supernatants were analyzed by focus forming assay.

### Focus forming assay

To measure HCV titer, supernatants from infected cells were serially diluted and then used to infect naïve Huh-7.5 cells in triplicate wells of a 48-well plate for 3 hours. At 48 hours post infection, cells were washed with PBS and fixed with 4% methanol-free paraformaldehyde (Sigma) for 30 minutes, and then washed again with PBS. Cells were then permeabilized (0.2% Triton-X-100 (Sigma) in PBS), blocked (10% FBS in PBS), and immunostained with a mouse anti-HCV NS5A antibody (1:500). Infected cells were visualized following incubation with horseradish peroxidase (HRP)-conjugated secondary mouse antibody (1:500; Jackson ImmunoResearch) and VIP Peroxidase Substrate Kit (Vector Laboratories). Foci were counted at 10X magnification, and viral titer was calculated using the following formula: (dilution factor × number of foci × 1000)/volume of infection (in μl), resulting in units of focus forming units / ml (FFU/ml).

### Immunoblotting

Cells were lysed in a modified RIPA buffer (10 mM Tris pH 7.5, 150 mM NaCl, 0.5% sodium deoxycholate, 1% Triton X-100) supplemented with protease inhibitor cocktail (Sigma) and phosphatase inhibitor cocktail (Millipore), and post-nuclear supernatants were harvested by centrifugation. Protein concentration was determined by Bradford assay, and 10 μg quantified protein was resolved by SDS/PAGE, transferred to either PVDF (for NS4A) or nitrocellulose membranes using either the Trans-Blot Turbo System (BioRad) or a wet system (BioRad), and blocked with either 3% bovine serum albumin (Sigma) in PBS with 0.1% Tween (PBS-T) or 10% FBS in PBS-T. Membranes were probed with specific antibodies against proteins of interest, washed 3X with PBS-T, and incubated with species-specific HRP-conjugated antibodies (Jackson ImmunoResearch, 1:5000), washed again 3X with PBS-T, and treated with Clarity enhanced chemiluminescence substrate (BioRad). Membranes were then imaged on X-ray film or by using a LICOR Odyssey FC. Immunoblots imaged using the LICOR Odyssey FC were quantified with ImageStudio software, and raw values of the protein of interest were normalized to those of controls (either Tubulin or GAPDH, as indicated). For immunoblots developed on film, Fiji was used (40). ImageStudio and Fiji give similar quantification results when compared directly.

### Immunoprecipitation

Quantified protein (between 80-160 μg) was incubated with protein-specific antibodies (either R anti-HA (Sigma) or anti-NS4A) in PBS at 4°C overnight with head over tail rotation. The lysate/antibody mixture was then incubated with either Protein A (for Flag-Riplet experiments) or Protein G Dynabeads (Invitrogen) for 2 hours. Beads were washed 3X in either PBS or RIPA for immunoprecipitation and eluted in 2X Laemmli Buffer (BioRad) supplemented with 5% 2-mercaptoethanol at 50°C for 5 minutes. Proteins were resolved by SDS/PAGE and subjected to immunoblotting as described above.

### Immunofluorescence analysis and confocal microscopy

Huh7 cells in 4-well chamber slides were fixed in 4% formaldehyde, permeabilized with 0.2% Triton-X-100, and immunostained with the following antibodies: mouse anti-HCV NS4A (Genotype 1B, 1:100, Virogen), rabbit anti-HA (1:100, Sigma), and rabbit anti-Sendai virus (SV) (1:1000, MBL International). Secondary antibody incubations were done with Alexa Fluor conjugated antibodies (Thermo Fisher) and with Hoescht (Thermo Fisher) for 1 hour. Following antibody incubations, slides were washed with 1X PBS, and mounted with ProLong Gold Antifade mounting medium (Invitrogen). Samples were imaged on a Zeiss 780 Upright Confocal using a 63X/1.25 oil objective and the 405, 488, 561, and 633 laser lines with pinholes set to 1 AU for each channel (Light Microscopy Core Facility, Duke University). Imaging analysis was done using Fiji software (40).

### Antibodies

Antibodies used for immunoblot and immunofluorescence analysis include: mouse anti-HCV NS4A (Genotype 1B, 1:1000, Virogen), mouse anti-HCV NS3 (Genotype 1B, 1:1000, Adipogen), mouse anti-HCV NS5A (Genotype 2A, 1:1000, clone 9e10, gift of Dr. Charles Rice, Rockefeller University), mouse anti-Tubulin (1:5000, Sigma), mouse anti-RIG-I (1:1000, Adipogen), anti-Flag-HRP (1:2500, Sigma), rabbit anti-Flag (1:1000, Sigma), rabbit anti-MAVS (1:1000, Bethyl Laboratories), mouse anti-IRF3 (1:1000, gift from Dr. Michael Gale Jr., University of Washington (41)), mouse anti-V5 (1:1000, Sigma), mouse anti-HA (1:1000, Sigma), rabbit anti-GAPDH (1:1000, Cell Signaling Technology), Hoescht (1:500, Thermo Fisher), Alexa Fluor conjugated secondary antibodies (1:500, Life Technologies), and rabbit anti-SV (1:1000, MBL International).

### IFN-β promoter luciferase assays

IFN-β promoter luciferase assays were performed by transfecting cells with pCMV-Renilla or pGL4.74 [hRluc/TK], pIFN-β-Luc, and expression plasmids as indicated. The following day, cells were infected with SV (Cantrell strain; Charles River labs). SV infections were performed in serum-free media at 200 hemagglutination units (HAU) for 1 h, after which complete media was replenished. At 20 hours post infection, cells were lysed, and a dual luciferase assay was performed (Promega). Values are displayed as relative luciferase units (RLU), which normalizes the Firefly luciferase (IFN-β-Luc) values to Renilla luciferase.

### Reverse transcription-quantitative PCR (RT-qPCR)

RNA was extracted from cell lysates using the RNeasy RNA extraction kit, and cDNA synthesis was performed on extracted RNA using iScript (BioRad). The resulting cDNA was diluted (either 1:3 or 1:4) in ddH_2_O. RT-qPCR analysis was performed using the Power SYBR Green PCR master mix (Thermo Fisher) on the QuantStudio 6 Flex RT-PCR system. The oligonucleotide sequences used for RT-qPCR are listed in Table 1. Heat map analysis was generated using Morpheus Software from the Broad (https://software.broadinstitute.org/morpheus). First the 2^ΔΔCt^ values (Comparative Ct Method) were calculated by setting the mock-infected Huh7 sample Ct value as the baseline for each biological replicate. Then, the mean of the SV-infected Huh7 triplicate samples is set to 1, and the relative fold induction for each gene between samples is shown.

### Statistical Analysis

Student’s unpaired *t* test or one-way ANOVA were implemented for statistical analysis of the data using GraphPad Prism software. Graphed values are presented as mean ± SD or SEM (n = 3 or as indicated); *p ≤ 0.05, **p ≤ 0.01, and ***p ≤ 0.005.

## Acknowledgements

We would like to thank the members of the Horner Lab and Dr. Heather Vincent for discussion and review of the manuscript, as well as Dr. Nicholas Barrows, Moonhee Park, Kevin Labagnara, Bianca Lupan, and Jason Willer for assistance with experiments, the Duke University Light Microscopy Core Facility, and the Duke Functional Genomics Core Facility. We also thank the following for reagents: Dr. Charles Rice at Rockefeller University, Dr. Michaela Gack at the University of Chicago, Dr. So Young Kim at the Duke University Functional Genomics Core, Dr. Michael Gale Jr. at the University of Washington, Dr. Adolfo Garcia-Sastre at Mount Sinai School of Medicine, and Dr. Shelton Bradrick and Dr. Mariano Garcia-Blanco at the University of Texas Medical Branch. This work was supported by funds from the National Institutes of Health (NIH): K22AI100935 (S.M.H.); R21AI124100 (S.M.H). Additional funding sources include the Burroughs Wellcome Fund (S.M.H); a Duke Bridge Award (S.M.H), and the Ford Foundation (CV).

